# Bactericidal Membrane Attack Complex formation initiates at the new pole of *E. coli*

**DOI:** 10.1101/2025.04.25.650575

**Authors:** Marije F.L. van ‘t Wout, Fabian Hauser, Philippa I.P. Holzapfel, Bart W. Bardoel, Carla J.C. de Haas, Jaroslaw Jacak, Suzan H.M. Rooijakkers, Dani A.C. Heesterbeek

**Affiliations:** Department of Medical Microbiology, University Medical Center Utrecht, Utrecht University, The Netherlands; University of Applied Sciences Upper Austria, Linz, Austria

## Abstract

Human immune protection against bacteria critically depends on activation of the complement system. The direct bacteriolytic activity of complement molecules against Gram-negative bacteria acts via the formation of Membrane Attack Complex (MAC) pores. Bactericidal MAC pores damage the bacterial outer membrane, leading to destabilization of the inner membrane. Although it is well-established that inner membrane damage is crucial for bacterial cell death, the critical event causing MAC-mediated inner membrane damage remains elusive. Here we questioned whether the bacterial cell envelope possesses vulnerable spots for MAC pores to insert. We followed the localization of MAC pores on *E. coli* over time using fluorescence microscopy and elucidate that MAC deposition initiates at the new bacterial pole, which induces inner membrane damage and halts bacterial division. MAC components C8 and C9 preferentially localize at new bacterial poles, while C3b localizes randomly on the bacterial surface. This suggests that preferential MAC localization is determined by one of the initial steps of MAC formation. These findings provide valuable information about the interplay between immune components and the Gram-negative cell envelope.

## Introduction

The immune system protects against bacterial infections by recognizing and attacking bacteria that invade the human body. One of the first lines of defense against pathogenic bacteria is the complement system, which consists of a large family of proteins in the blood and other body fluids (1). The importance of this system in the defense against bacteria is illustrated by the recurrent infections that occur in people with complement deficiencies (2,3) or patients treated with complement inhibitors (4). Complement proteins can become activated upon recognition of bacteria, starting a cascade of cleavage reactions on the bacterial surface. Complement activation triggers multiple effector mechanisms that aid in bacterial clearance, one of them being the formation of Membrane Attack Complex (MAC) pores. MAC pores are formed when C5 convertase enzymes on the bacterial surface generate C5b, which can assemble with C6, C7, C8 and up to 18 copies of C9 to form a complete MAC pore (5). By causing large-scale disruption of the bacterial cell envelope, MAC pores can directly kill Gram-negative bacteria (6,7).

Gram-negative bacteria possess a complex cell envelope including both an inner and outer membrane and a layer of peptidoglycan in between. The outer leaflet of the outer membrane is primarily composed of lipopolysaccharides (LPS), but also contains outer membrane proteins (OMPs) (8). The bacterial cell envelope exhibits some degree of spatial heterogeneity, as peptidoglycan synthesis and OMP insertion primarily take place at the division poles (9). Due to this process, older OMPs and peptidoglycan shift towards the old poles, where mature peptidoglycan inhibits OMP insertion (9). Despite these spatial differences within the bacterial cell envelope, its impact on complement deposition and MAC-mediated killing has not yet been investigated.

Stable insertion of MAC pores into the bacterial outer membrane eventually results in inner membrane damage, which is the crucial event for bacterial cell death (7,10,11). Although it is hypothesized that inner membrane damage is an indirect effect of outer membrane damage, the critical event for MAC-mediated inner membrane damage remains unknown. Structural imaging studies revealed that MAC pores assembled from purified complement components C5b6 and C7-C9 form an asymmetric pore that is able to rupture single membrane particles (12–14). MAC-mediated killing of Gram-negative bacteria requires a more complicated process due to the complex structure of the bacterial cell envelope. In order to properly insert into the bacterial outer membrane and damage the bacterial inner membrane, MAC pores need to form close to the bacterial surface by convertase enzymes (10). Atomic Force Microscopy (AFM) on *E. coli* revealed that these bacteria can become fully covered with convertase-generated MAC pores at high complement concentrations and after sufficient incubation times (7). It is however unknown where MAC pores localize at the moment inner membrane damage is triggered, and whether some locations within the bacterial cell envelope are more vulnerable to MAC insertion.

Here we visualized MAC localization on *E. coli* over time and reveal that initial MAC deposition occurs with a distinct localization pattern. By simultaneously capturing the moment of inner membrane damage, we show that MAC insertion at new bacterial poles coincides with bacterial inner membrane damage. These polar MAC pores also severely impair bacterial growth and division. Altogether, these findings shed light on how complement molecules attack the complex cell envelope of Gram-negative bacteria.

## Results

### MAC pores that trigger inner membrane damage are deposited with a distinct localization pattern

First, we wanted to investigate whether bactericidal MAC pores preferentially localize at certain parts of the bacterial surface. Therefore, we analyzed MAC deposition on *E. coli* MG1655 over time and captured the moment where the number of MAC pores was just sufficient to cause bacterial killing. To solely look at the effect of MAC on bacterial killing in the absence of other serum components, we pre-labeled bacteria with C5 convertase enzymes by incubating them with C5-depleted serum (10). Then, after washing, convertase-labeled bacteria were incubated with purified MAC components (C5-C9) at a concentration of 2.5% serum equivalent (15) for different amounts of time (Fig. 1A). MAC deposition was visualized by using C9 that was fluorescently labeled with AF647. Previously, we showed that inner membrane damage is a critical event for MAC-mediated bacterial killing (10,11). Therefore, to measure at which timepoint MAC pores were able to induce bacterial cell death, we stained for inner membrane damage with a naturally membrane impermeable DNA dye (Sytox Blue).

**Figure 1:**
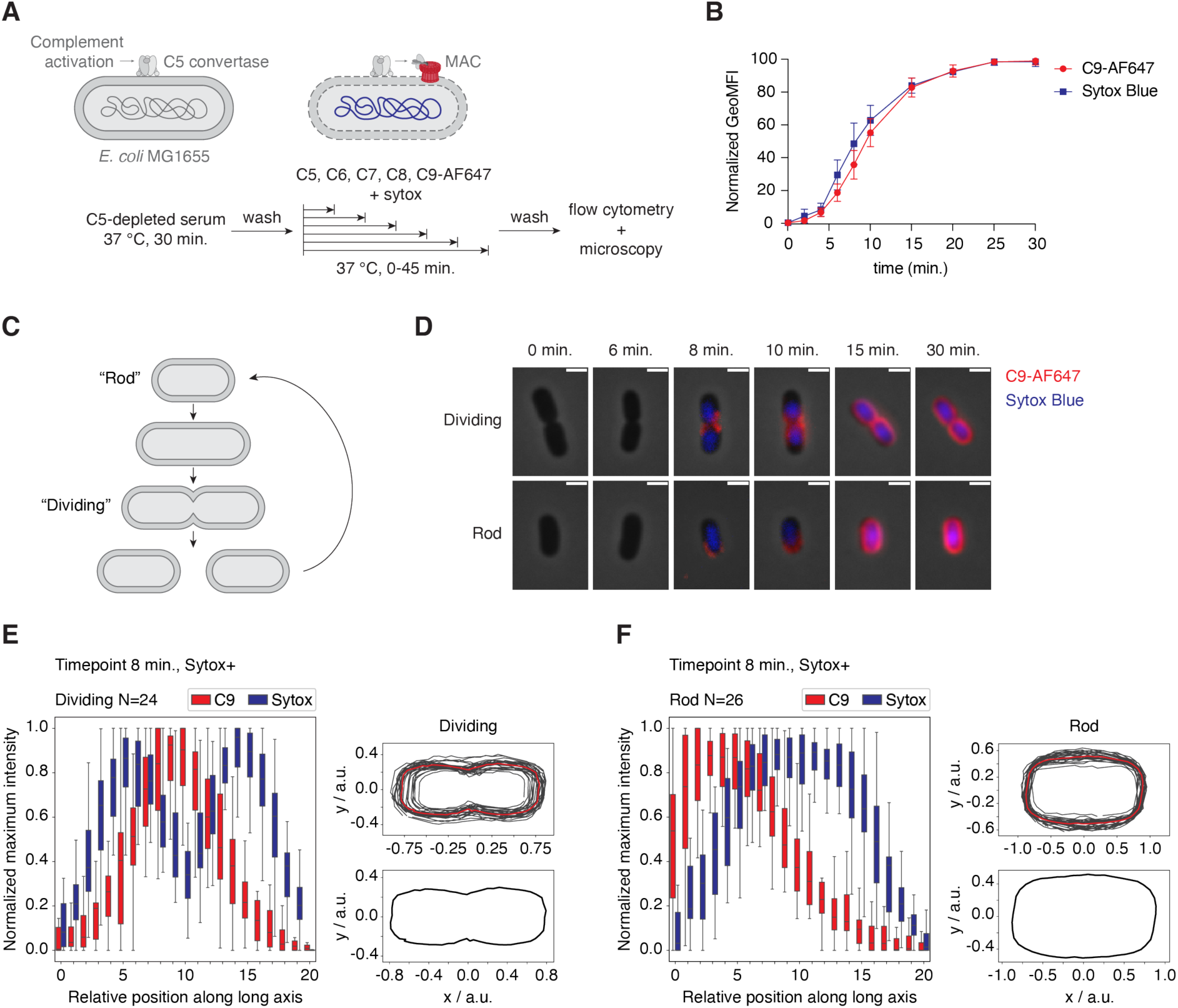
MAC pores that trigger inner membrane damage are deposited with a distinct localization pattern. (**A**) Schematic overview of the experimental setup used to study MAC deposition and inner membrane in time. E. coli MG1655 was pre-incubated with C5-depleted serum, washed, and then incubated with purified MAC components for different amounts of time. (**B**) C9-AF647 deposition and inner membrane damage (Sytox Blue) in time, measured by flow cytometry. GeoMFI values were normalized by calculating the percentage of the maximum value after log-transformation. Data represent mean ± SD of four independent experiments. (**C**) Schematic overview of the different stages that occur during bacterial growth and division. (**D**) Microscopy analysis of the samples in (B). C9-AF647 is shown in red and Sytox in blue. Individual bacteria reflect an average bacterium in the sample and are representative for three independent experiments. Scale bars, 2 µm. (**E**) On the left, normalized average C9 and Sytox distribution along the long axis of dividing bacteria after 8 minutes incubation with C5-C9. On the right, the average shape of bacteria that were classified as dividing. Data analysis was performed on multiple images of one independent experiment. (**F**) On the left, normalized average C9 and Sytox distribution along the long axis of rod-shaped bacteria after 8 minutes incubation with C5-C9. On the right, the average shape of bacteria that were classified as rod-shaped. Data analysis was performed on five images of one independent experiment.

By using flow cytometry, we first confirmed that MAC deposition increased gradually over time and that these pores could indeed induce inner membrane damage (Fig. 1B, Fig. EV1A, B). To visualize where MAC pores localize at the moment inner membrane damage is triggered, we imaged the samples that were exposed to MAC components for different amounts of time by widefield fluorescence and phase-contrast microscopy. Since bacteria were in different stages of the cell cycle within one sample, we categorized them into two groups: ‘rod-shaped’ and ‘dividing’ (Fig 1C). On bacteria with a clear septum at mid-cell (dividing), MAC deposition initiated at the division poles (Fig. 1D). On bacteria that were just formed after cell separation (rod-shaped), MAC pores localized on only one of the two outer poles (Fig. 1D). On both groups of bacteria, polar MAC deposition was sufficient to cause inner membrane damage. MAC deposition increased over time until bacteria were completely covered with MAC pores, consistent with previous studies (7). To show that this localization pattern is representative for other bacteria within the sample, we developed a high-throughput image analysis pipeline to identify individual bacteria within our images, categorize them into different growth stages and determine the average fluorescence intensity profile along the long axis of each cell (Fig. EV2). This image analysis pipeline can be applied to large data sets and therefore allowed us to quantify average fluorescence distributions on multiple bacteria. To prove that the algorithm works correctly, it was tested on simulations of rod-shaped and dividing bacteria that were fully covered with fluorescent signal and showed random fluorescence distribution (Fig. EV3). By applying the pipeline to images of a timepoint at which we first start to observe bacteria with inner membrane damage (8-minute timepoint), we demonstrate that the specific MAC localization shown in Fig. 1D is representative for other bacteria in the sample (Fig 1E, F). By orienting bacteria based on the C9-AF647 signal, having the side with the highest fluorescence intensity on the left, we confirm that MAC pores preferentially localize at mid-cell on dividing bacteria, and on one of the outer poles of rod-shaped bacteria. To show that this localization pattern can be observed on more *E. coli* strains, we performed the same experiment on a MAC-sensitive clinical *E. coli* isolate, EC10. We found the same localization pattern of MAC pores on EC10, also correlating with bacterial inner membrane damage (Fig. EV4).

Complement activation can occur through three different pathways: the classical pathway, triggered by antibody-antigen binding; the lectin pathway, triggered by mannose-binding lectin or ficolin binding; and the alternative pathway, triggered by spontaneous C3 hydrolysis, but also acting as an amplification loop (1). Since pre-incubation of *E. coli* in C5-depleted serum could result in complement activation through each of these routes, we investigated whether only antibody-dependent complement activation triggers the same preferential MAC localization. Therefore, we performed a similar analysis in a fully purified complement assay in which only the classical pathway is activated, as all complement components necessary for classical pathway activation are added individually (Fig. 2A) (15). To activate complement, we used an antibody that recognizes GlcNAc moieties present in the LPS of *E. coli* (*Muts et al.* (*2025), manuscript under review*). At the antibody concentration that was used in our experimental setup, bacteria were fully covered with IgM without showing preferential localization (Fig. EV5). Also in this experimental setup, MAC deposition and inner membrane damage increased gradually over time (Fig. 2B), but started earlier than in the serum setup (Fig. 1B). Similar to the serum setup, MAC localized at the division poles on dividing bacteria and at one of the two outer poles on rod-shaped bacteria at the moment inner membrane damage was first induced (Fig. 2C-E). Together, these results show that bactericidal MAC pores have a distinct localization pattern on *E. coli*.

**Figure 2:**
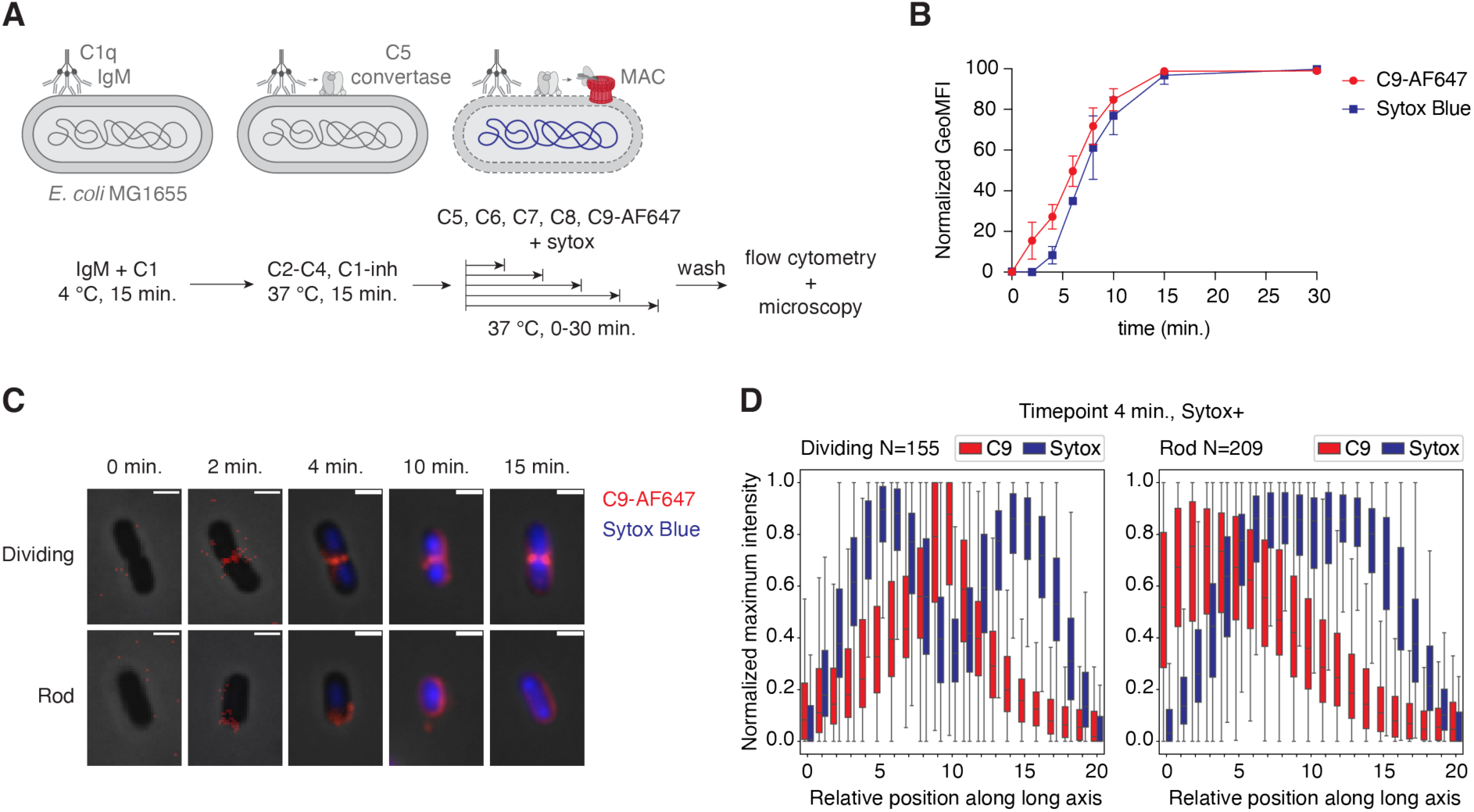
Specific localization of MAC pores after antibody-mediated complement activation. (**A**) Schematic overview of the purified classical pathway setup used to study MAC localization and inner membrane in time. (**B**) C9-AF647 deposition and inner membrane damage (Sytox Blue) in time, measured by flow cytometry. GeoMFI values were normalized by calculating the percentage of the maximum value after log-transformation. Data represent mean ± SD of three independent experiments. (**C**) Phase-contrast and overlayed fluorescence images of selected bacteria at different timepoints. C9-AF647 is shown in red and Sytox in blue. Individual bacteria have been chosen to reflect an average bacterium in the sample and are representative for three independent experiments. Scale bars, 2 µm. (**D**) Normalized average C9 and Sytox distribution along the long axis of dividing and rod-shaped bacteria after 4 minutes incubation with C5-C9. Data analysis was performed on 19 images of one independent experiment.

### MAC pores preferentially deposit at the new pole of *E. coli*

Next, we investigated whether MAC pores preferentially localize at a specific pole on rod-shaped bacteria. Bacterial poles can be classified as old or new, the latter being derived from the division pole in the previous round of division (Fig. 3A). Given that MAC deposits at the division poles of dividing bacteria, we hypothesized that it localizes at the new pole of rod-shaped bacteria. To distinguish the old and new poles, we used an approach to fluorescently label the old poles of *E. coli* with HCC-amino-D-alanine (HADA). HADA is a fluorescent D-alanine which is incorporated into newly synthesized peptidoglycan. By first adding HADA to the growth medium of *E. coli* for one hour, we ensured that the peptidoglycan became uniformly labeled (Fig 3A, B and (16)). After washing, we allowed the bacteria to grow in the absence of HADA for another 45 minutes (Fig. 3B). Since peptidoglycan is primarily synthesized at the bacterial septum, the formation of new, unlabeled peptidoglycan causes HADA-labeled peptidoglycan to move towards the old poles (Fig. 3A and (16)). During this process, bacteria were simultaneously labeled with convertases and exposed to purified MAC components (Fig. 3B), similar to the setup described in figure 1A. By using this method, we show that MAC pores localize at the poles that are not labeled with HADA, indicating that they specifically localize at new bacterial poles (Fig. 3C). Analysis of a dataset of >3900 individual bacteria was used to validate the localization of MAC pores at the new pole of rod-shaped *E. coli* (Fig. 3D). For dividing bacteria, the HADA signal was mostly oriented towards the old pole of one of the two daughter cells, with MAC pores again localizing at the division poles (Fig. 3D). This confirms that MAC pores preferentially localize at the new poles of both dividing and rod-shaped bacteria.

**Figure 3:**
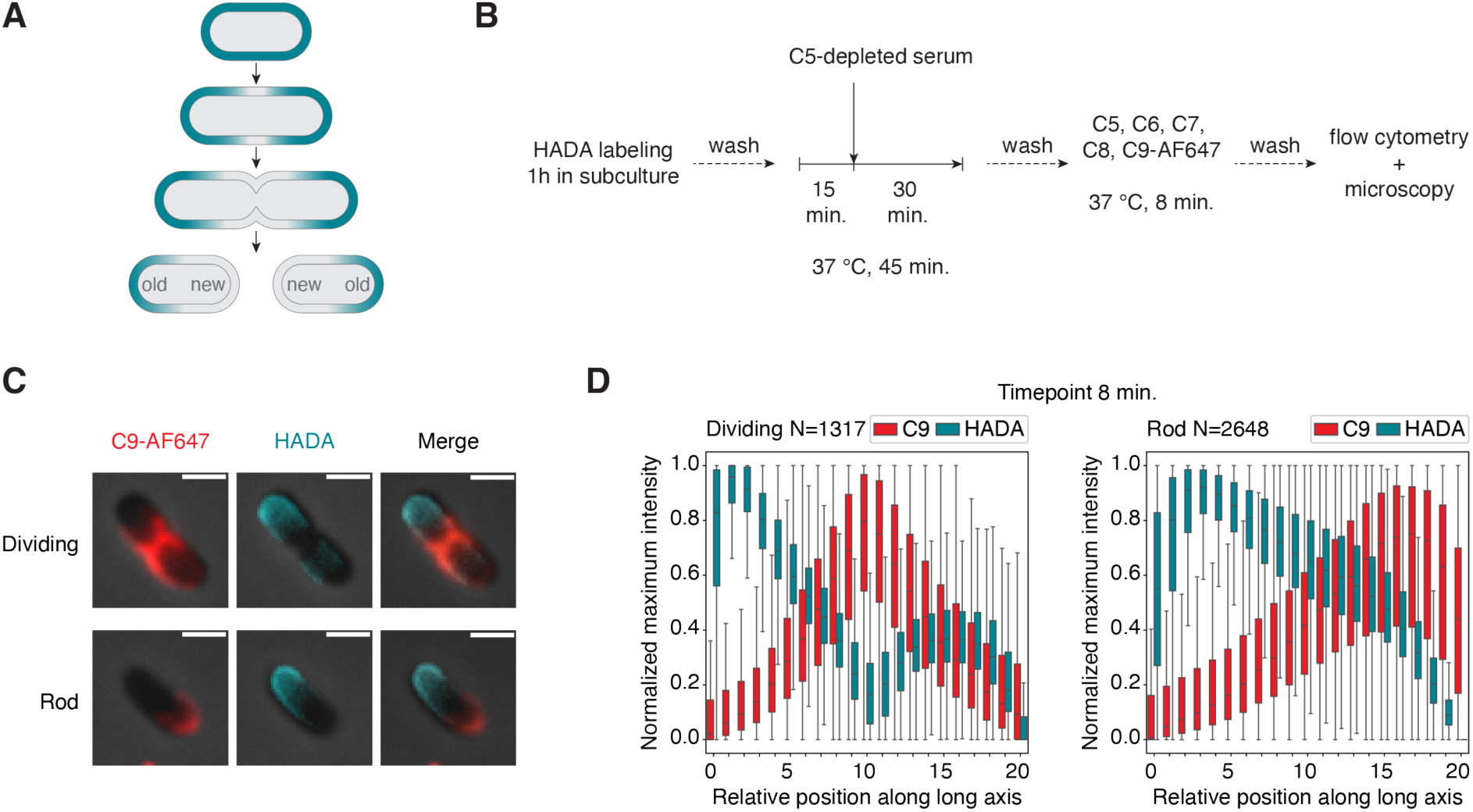
MAC pores specifically localize at the new pole of E. coli. (**A**) Schematic representation of HADA distribution (in cyan) upon bacterial growth. After labeling bacterial peptidoglycan with HADA, new unlabeled peptidoglycan is synthesized at the bacterial septum, moving labeled peptidoglycan toward the old poles (**B**) Schematic overview of the experimental setup used to label old bacterial poles with HADA and induce MAC deposition. After labeling the peptidoglycan of E. coli MG1655 with HADA, bacteria were allowed to grow in absence of HADA for 45 minutes. Simultaneously, bacteria were incubated with C5-depleted serum, washed, and then incubated with purified MAC components for 8 minutes. (**C**) Phase-contrast and overlayed fluorescence images of selected bacteria after performing the assay as shown in (B). C9-AF647 is shown in red and HADA in cyan. Individual bacteria have been chosen to reflect an average bacterium in the sample and are representative for three independent experiments. Scale bars, 2 µm. (**D**) Normalized average C9-AF647 and HADA distribution along the long axis of dividing and rod-shaped bacteria. Data analysis was performed on 50 images of one independent experiment.

### MAC pores at the new pole of *E. coli* impair bacterial cell division

Next, we analyzed how bacterial growth is affected by the deposition of MAC pores at the new pole. We have previously shown that inner membrane damage correlates with a reduction of colony forming units (CFUs) (10,11). However, CFU counts only provide information about the percentage of bacteria that survives and does not show whether individual bacteria can still grow. To study how MAC localization affects bacterial growth and division, we used time-lapse microscopy to follow individual bacteria for 2.5 hours after exposure to MAC components. First, we confirmed that also in this experimental setup (Fig. 1A), an increase in inner membrane damage correlated with a decrease in CFUs (Fig. 4A). In line with the partial survival of bacteria after 6, 8 and 10 minutes of exposure to MAC components, the number of cell divisions per bacterium differed vastly within one sample (Fig. 4B and Movies EV1-4). To study whether the number of divisions was dependent on the amount of MAC pores per bacterium, we manually categorized bacteria based on inner membrane damage (Sytox-positivity) and the level of MAC deposition (Fig. EV6). Bacteria were first classified as either Sytox-negative or -positive. Sytox-positive were further categorized into having polar C9 or being fully covered with C9. This categorization revealed that differences in bacterial growth could largely be explained by inner membrane damage, as bacteria with inner membrane damage showed drastically impaired cell division (Fig. 4C). Furthermore, most bacteria with MAC pores at the new pole were unable to grow and divide (Fig. 4C, D). Interestingly, in some cases, bacteria with inner membrane damage and polar MAC pores were able to partially escape killing (Fig. 4D). This suggests that some bacteria can survive a low level of inner membrane damage. Bacteria with an intact inner membrane at the start of the video were able to go through an average of ∼4 divisions (Fig. 4D). There was a large variation in the number of divisions per bacterium, but this was also seen for bacteria that had not been exposed to MAC pores (0-minute timepoint) (Fig. 4D). When bacteria were exposed to MAC components for 10 minutes, almost all imaged bacteria were fully covered with MAC pores and unable to divide (Fig. 4C, D). This correlates with almost no bacterial survival (CFUs) after 10 minutes of exposure to MAC components (Fig. 4A). Taken together, this shows that MAC pores at the new pole of *E. coli* severely impair bacterial cell growth and division.

**Figure 4:**
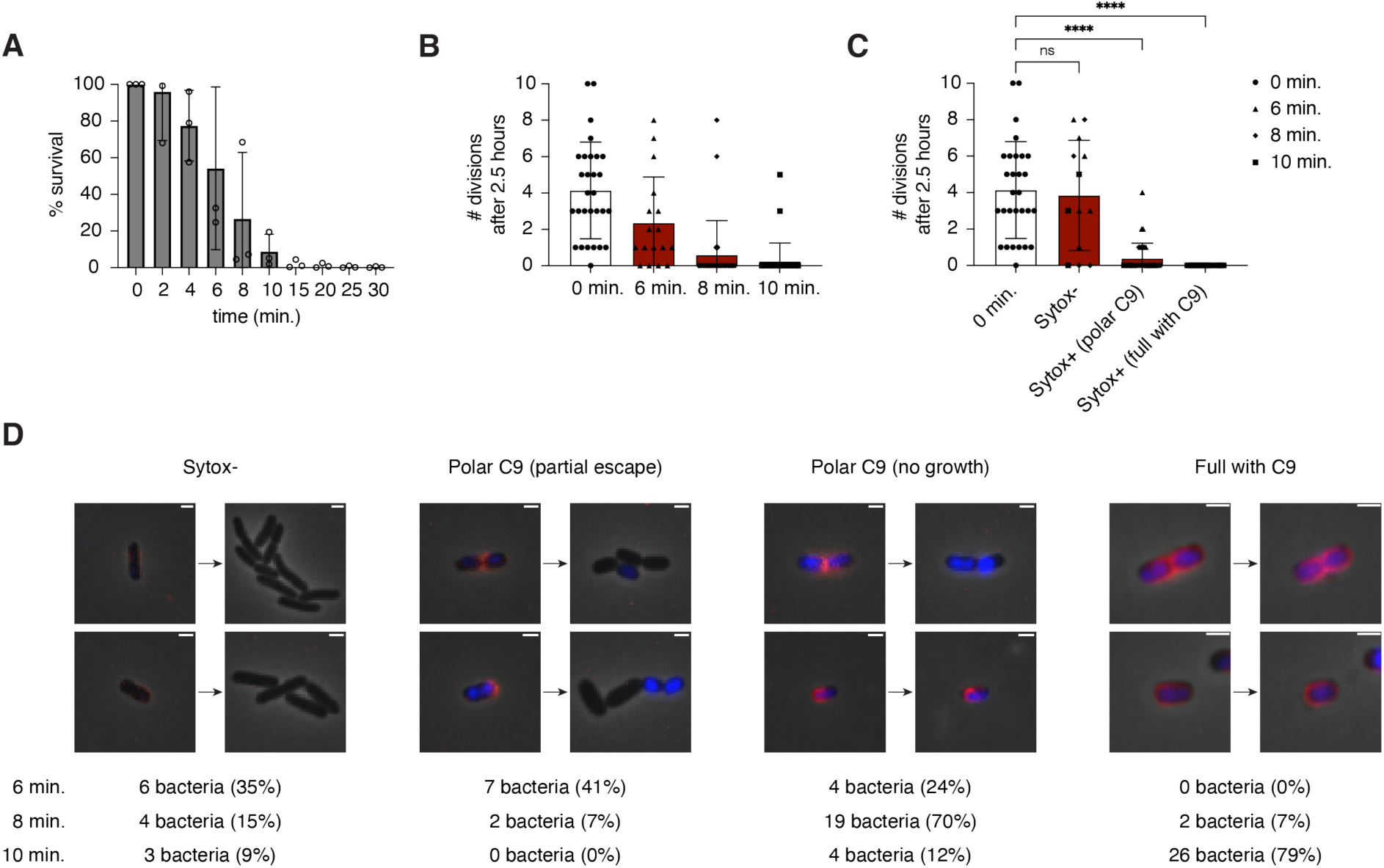
MAC pores at the new pole of E. coli drastically impair bacterial cell division. E. coli MG1655 was pre-incubated with C5-depleted serum, washed, and then incubated with purified MAC components for different amounts of time. (**A**) Bacterial survival after exposure to MAC components based on CFU/ml. Percentage survival was calculated by normalizing to the 0-minute timepoint, in which bacteria had not been exposed to MAC components. Data represent individual values with mean ± SD of three independent experiments. (**B**) The number of divisions of individual bacteria within 2.5 hours, imaged after exposure to MAC components for the indicated amounts of time. Data represent individual values with mean ± SD. (**C**) Graph as in (B), grouping bacteria exposed to MAC components based on Sytox-positivity and the localization of C9-AF647. Data represent individual values with mean ± SD. Statistics was performed using a Kruskal-Wallis test followed by a Dunn’s post-hoc test, comparing each group with the 0-minute timepoint. (**D**) Images of bacteria that had been incubated with C5-C9 for 6-10 minutes, before and after allowing growth for 2.5 hours. Images show examples of the groups shown in (C). Below is the number and percentage of bacteria showing each phenotype after 6, 8 or 10 minutes incubation with C5-C9. Scale bars, 2 µm.

### Deposition of MAC component C8 also starts at the new pole of *E. coli*

Next, we questioned what step in the complement cascade determines the preferential localization of C9 at the new pole. To study whether the C5b-8 complex also shows a distinct localization pattern or whether C9 localization may be caused by more efficient C9 polymerization at the new pole, we simultaneously measured C8 and C9 deposition. For this, we exposed *E. coli* to MAC pores for 8 minutes using the same setup as described in figure 1 but using recombinant C8 that was directly labeled with AF488. At this timepoint, we observed co-localization between C8-AF488 and C9-AF647 at the bacterial (division) poles (Fig. 5A, B). Since C9 might cause more efficient anchoring of C8 at the new pole, we next studied the deposition of C8 molecules in the absence of C9. By exposing convertase-labeled bacteria to C5-C7 and C8-AF488 for 8 minutes, we show that C8 preferentially inserts at the bacterial poles independently of C9 (Fig. 5C, D). Finally, we wanted to know whether more efficient complement activation at the new pole could explain the localized insertion of C8 and C9. Therefore, we measured the distribution of C3b molecules which are essential components of the C5 convertase enzymes. C5 convertases trigger MAC formation by cleaving C5, thereby forming the first MAC component C5b. C3b molecules were evenly distributed over the bacterial surface after incubation in C5-depleted serum (Fig. 5E, F), indicating that the preferred polar insertion of C8 and C9 is independent of C3b. Altogether, this suggests that the preferential localization of MAC pores at the new pole of *E. coli* is determined by one of the initial steps of MAC formation.

**Figure 5:**
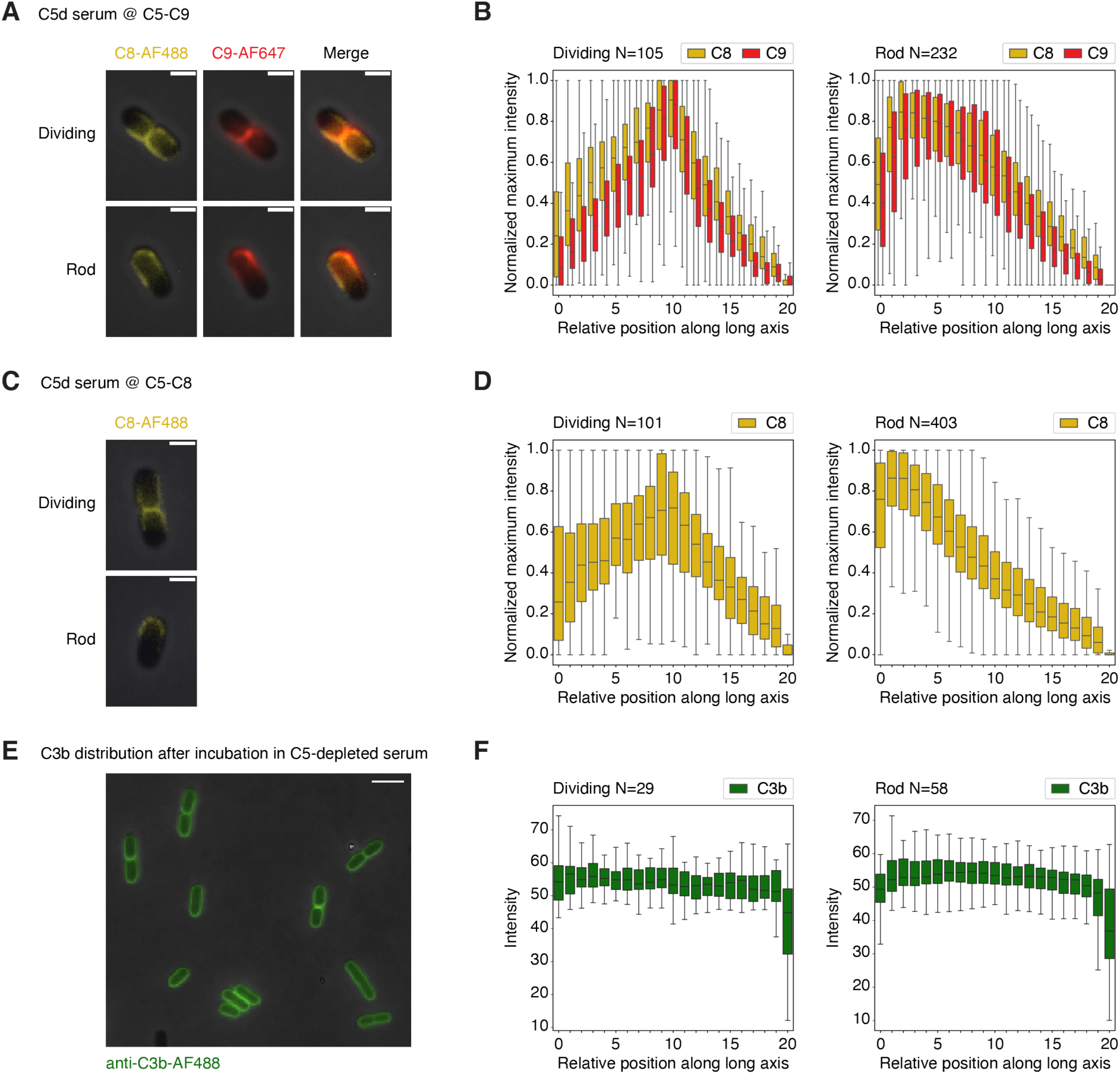
Deposition of MAC component C8 also starts at the new pole of E. coli. (**A**) Convertase-labeled E. coli bacteria were exposed to C5-C7, C8-AF488 and C9-AF647 for 8 minutes. Localization of C8-AF488 (yellow) and C9-AF647 (red) was determined by phase-contrast and widefield fluorescence microscopy. Scale bars, 2 µm. (**B**) Data analysis of the sample shown in (A). Graphs show the normalized average C8-AF488 and C9-AF647 distribution along the long axis of dividing and rod-shaped bacteria. Data analysis was performed on 20 images of one independent experiment. (**C**) Complement deposition on E. coli was induced as in (A), but in the absence of C9. Localization of C8-AF488 (yellow) was determined by phase-contrast and widefield fluorescence microscopy. Scale bars, 2 µm. (**D**) Data analysis of the sample shown in (C). Graphs show the normalized average C8-AF488 distribution along the long axis of dividing and rod-shaped bacteria. Data analysis was performed on 20 images of one independent experiment. (**E**) Bacteria were incubated with 10% C5-depleted serum followed by staining for C3b using a fluorescently labeled monoclonal antibody. Scale bar, 5 µm. (**F**) Data analysis of the sample shown in (E). Graphs show the average anti-C3b-AF488 distribution along the long axis of dividing and rod-shaped bacteria. Data analysis was performed on three images of one independent experiment. All images and graphs are representative for three independent experiments.

## Discussion

In this study, we reveal that bactericidal MAC pores show a distinct localization pattern on *E. coli.* By capturing the start of MAC deposition, we show that MAC pores are first deposited at new bacterial poles, where they trigger inner membrane damage and halt bacterial cell division. Whereas MAC pores initially insert at the new pole, we found that bacteria become fully covered with MAC pores within approximately 15 minutes of exposure to MAC components. This is in line with a recent study in which MAC-treated *E. coli* was imaged by AFM, which showed random distribution of MAC pores after these exposure times (7). Since initial MAC deposition at the new pole turned out to be sufficient to trigger bacterial cell death, the question remains how relevant later-formed pores at different locations are in bacterial killing. We previously showed that a limited number of C8 molecules (roughly between 100-1000 per bacterium), and thus MAC pores, are sufficient to cause inner membrane damage (10). Future studies should point out whether MAC insertion at the new pole is a requirement for bacterial cell death, or whether MAC pores at other locations can also efficiently induce bacterial killing. If MAC pores at other locations can also induce inner membrane damage, the here-observed preferential localization of MAC pores is a matter of insertion efficiency at these places. A high local density of MAC pores at the new pole resulting from more efficient insertion may also contribute to causing local large-scale disruption of the outer membrane. The observation that MAC deposition initiates at new bacterial poles, which are formed during cell division, poses the question whether active division is a requirement for MAC-dependent killing of Gram-negative bacteria.

In order to understand how the complement system attacks bacterial cells, it is essential to consider the impact of the complex structural organization of the bacterial cell envelope. The pattern of MAC localization seems to resemble images of the coordinated assembly of peptidoglycan and insertion of OMPs into the outer membrane of *E. coli* (9). The authors revealed that upon cell division, bacteria preferentially insert OMPs at the division poles, depending on the synthesis of immature peptidoglycan at the same locations. Preferential insertion of MAC pores at these exact spots could indicate that the assembly of cell envelope components creates a vulnerable spot within the cell envelope during bacterial division. This vulnerability may not be unique to the insertion of MAC pores but may also apply to other membrane-attacking compounds. For example, the antimicrobial peptide Cecropin A was shown to preferentially attack septating cells and cause leakage of periplasmic GFP initially from the division poles (17). This supports the hypothesis that bacterial cell division may coincide with increased vulnerability to compounds that attack the bacterial membrane.

Time-lapse microscopy of bacterial growth after MAC deposition demonstrated that polar MAC deposition and inner membrane damage correlate with bacterial cell death. Due to heterogeneity in the exact timing of MAC deposition, some bacteria within a sample are already killed while others are not yet. When categorizing bacteria based on the localization of the MAC, we found that MAC deposition at new bacterial poles was sufficient to cause inner membrane damage and halt bacterial growth. This again proves that relatively low amounts of MAC pores are sufficient to induce bacterial killing. Interestingly, in some cases we noticed that bacteria with limited MAC deposition, but detectable inner membrane damage were able to survive and divide again. This partial escape of bacterial killing, which also caused a loss of Sytox signal over time, suggests that bacteria can repair low degrees of inner membrane damage. However, MAC deposition at the bacterial poles was often sufficient to stop bacterial growth entirely.

The exact step within the complement cascade that causes preferred polar MAC deposition remains to be determined. We show that MAC component C8 has the same localization pattern as C9, indicating that it is not C9 polymerization that is more efficient at the new pole. Furthermore, we observed homogeneous distribution of the IgM antibody targeting GlcNAc moieties present in the LPS core of *E. coli*. This indicates that this antibody does not preferentially localize at certain locations, which would have caused more efficient complement activation at these spots. This is further supported by the random distribution of C3b, an essential component of C5 convertases, again suggesting that polar MAC localization is not caused by more potent complement activation at the new pole. While more efficient insertion of MAC precursors into potentially weak bacterial poles seems a likely explanation for the preferred polar MAC deposition, it could also be caused by enhanced activity of C5 convertases at these spots. For instance, even though C3b seems to be uniformly distributed over the bacterial surface, there might be a subtle difference in the local density of C3b at the new bacterial pole. Follow-up studies are needed to study whether preferential MAC localization at the new poles may be caused by more efficient C5 conversion at these locations.

Our observation of polar MAC localization on the MAC-sensitive *E. coli* MG1655 strain were validated on a MAC-sensitive *E. coli* clinical isolate. Future studies are needed to investigate whether these findings can also be translated to other Gram-negative bacterial species. To our knowledge, specific localization of MAC pores on Gram-negative bacteria has not yet been described before. In contrast, the localization of MAC pores has previously been tested on four different Gram-positive bacterial species and varied from being predominantly present at the division pole, to a more random distribution (18). Although Gram-positive bacteria possess a vastly different cell envelope and are intrinsically MAC-resistant, it may be interesting to include these and other Gram-negative species with different sensitivities to MAC-dependent killing in follow-up studies. Extending our current analysis to a larger bacterial panel may guide us towards a better understanding of what determines MAC localization and how this relates to MAC sensitivity. A relatively large proportion of pathogenic bacteria is resistant to MAC-mediated killing, which is often linked to improper membrane insertion of MAC pores (19). On Gram-negative bacteria, the insertion of MAC pores is likely influenced by the presence of an O-antigen (20,21) or polysaccharide capsule (22,23). Since the laboratory *E. coli* strain used in this study lacks an O-antigen and capsule, testing MAC localization on Gram-negative bacteria with varying O-antigen lengths and capsule thicknesses may reveal whether these structures influence the distribution of MAC pores on the bacterial surface. If MAC insertion at the new pole turns out to be required for MAC-mediated killing, bacteria might have acquired defense mechanisms against MAC-dependent killing by preventing MAC insertion at this location.

Taken together, this study provides novel insights into how Gram-negative bacteria are killed by MAC pores. The growing problems with antibiotic resistant bacteria (24) have increased the urgency to invest in the development of alternative antimicrobial therapies (25–27). These for example include the use of antibodies to activate the complement system and trigger MAC formation, the discovery of new antimicrobial agents that directly act on Gram-negative bacteria or the use of lytic bacteriophages. The here-presented fundamental insights into what locations of the bacterial cell envelope may be most vulnerable to components that act on the bacterial outer membrane could provide valuable information in the design of these new antibacterial therapies.

## Acknowledgements

The work was funded by the European Research Council (ERC) under the European Union’s Horizon 2020 research and innovation program (grant agreement no. 101001937, ERC-AC-CENT, to SHMR) and by the Austrian Forschungsförderungsgesellschaft (FFG, Nano-Carriers, project number: 898921). We would like to acknowledge Marek Basler, Johannes Schneider and Andrea Vettiger for the valuable technical input on live-cell imaging of bacteria and Jos van Strijp for proofreading of the manuscript. Furthermore, we would like to thank Floor Annabel Niessen for providing us with advice on statistical tests used.

## Author contributions

MFLW, FH, BWB, JJ, SHMR and DACH designed research; MFLW, PIPH, CJCH and DACH performed experiments; FH and JJ designed a pipeline for automated image analysis, MFLW, FH, PIPH and DACH analyzed the data and MFLW, FH, BWB, JJ, SHMR and DACH wrote the paper. JJ, SHMR and DACH supervised the project. JJ and SHMR acquired funding. All authors approved the final version of the manuscript.

## Disclosure and competing interests statement

The authors declare that they have no conflict of interest.

## Methods

### Bacterial strains

Unless specified in the figures, experiments were performed on *E. coli* strain MG1655. The *E. coli* clinical isolate (EC10) was isolated from a patient with bacteremia and obtained from the Medial Microbiology department of the University Medical Center Utrecht. For each experiment, a single bacterial colony was picked from a blood agar plate and grown overnight in LB medium, shaking at 37 °C. The next day, cultures were diluted 1:100 in LB medium and grown until mid-log phase (OD600nm = 0.35-0.65).

### Reagents

C5-depleted serum and complement proteins C1, C4, C8 and C1-inhibitor were obtained from Complement Technology. Production and fluorescent labeling of C8-AF488 and C9-AF647 is described below. His-tagged C2, C5, C6, and C7 were expressed in Expi293F cells and purified using the same protocol as for C8 and C9. C3 was isolated from human plasma as described previously (28). Monoclonal anti-GlcNAc (4497) IgM was produced as described previously (*Muts et al.* (*2025), Manuscript under review*). Sytox Blue was purchased from ThermoFisher. Concentrations of complement components in 100% serum: 135 µg/ml C1, 20 µg/ml C2, 1250 µg/ml C3, 400 µg/ml C4, 70 µg/ml C5, 64 µg/ml C6, 56 µg/ml C7, 55 µg/ml C8, 60 µg/ml C9, 180 µg/ml C1-inhibitor. Unless stated otherwise, incubations were performed in RPMI 1640 medium supplemented with 0.05% human serum albumin (RPMI/HSA).

### Fluorescent labeling of complement component C9

C9-AF647 was produced by modifying the C-terminus of C9 (NM_001737;AA22-559) with a C-terminal 3xLinker (3xGGGGS)-LPETGG-6xHIS tag which was expressed in EXPI293F cells, as described in (29). After purification, 50 μM C9-3xGGGGS-LPETG-6xHIS was incubated for 2h at 4 °C with 500 μM ggg-AF647 (Click Chemistry Tools) and 10 μM His-TEV-Sortase 7+ in 10 mM CaCl_2_, 50 mM Tris, 150 mM NaCl; pH 8.0 buffer. Then, 0.1 volume of 10x binding buffer (50 mM Tris/3.5 M NaCl/200 mM imidazole; pH8.0) was added. To capture the His-TEV-Sortase and non-AF647 labeled (His-tag labeled) C9, the sample was loaded onto a HisTrap FF column (GE Healthcare). The flow through (C9-AF647) was collected, concentrated to 500 µl, and subsequently applied to a Superdex 200 increase 10/300 GL (Cytiva) gel filtration column. Fractions containing AF647-labeled C9 were collected and analyzed to have a AF647 dye to protein label (DOL) of 56%.

### Fluorescent labeling of complement component C8

The genes for C8α (NM_000562), C8β (NM_000066) and C8y (NM_000606) were cloned in the pcDNA3.4 TOPO vector (Thermo Fisher Scientific) and expressed in EXPI293F cells, as described in (29). In short, the cystatin(S) signal peptide was used for all 3 genes. C8α (AA31-584) was cloned to express a C-terminal 3xLinker (3xGGGGS)-LPETGG-StrepTagII (SAWSHPQFEK). Both C8β (AA55-591) and C8y (AA21-202) were cloned to express a C-terminal 3xLinker (3xGGGGS)-LPETGG-6xHIS tag. The LPETG sequence in the tag of each C8 subunit was used for later triple labeling of C8 via sortagging (30). Expression in EXPI293F cells was performed using a total of 0.5 µg plasmid/ml cells with a ratio of C8α:C8β:C8y:Empty vector of 0.24:0.33:0.24:0.19 per µg total used plasmid. After 5 days of expression, EXPI293F supernatant was buffer exchanged to 100 mM Tris, 150 mM NaCl, 1 mM EDTA; pH8.0 using the QuixStand benchtop system (GE HealthCare). To inactivate residual biotin, 1.8 µl/ml Biolock (IBA Lifesciences) was added and incubated for 30 minutes. Then, the C8 supernatant was applied to a 1 ml Strep-Tactin^®^XT 4Flow^®^ FPLC column (IBA Lifesciences) and C8 was eluted according to the manufacturer’s description. For triple labeling of C8, 4.5 µM C8 was treated for 2h at 4°C with 45 µM ggg-AF488 (Thermo Fisher Scientific), 10 mM CaCl_2_, 0.9 µM HIS-TEV-Sortase 7+ (31) in 50 mM Tris, 150 mM NaCl; pH8.0 buffer. Then, 0.1 volume of 10x Strep-Tactin buffer (500 mM Tris/ 150 mM NaCl/110 mM EDTA; pH8.0) was added, before application to a 1 ml Strep-Tactin^®^XT 4Flow^®^ FPLC column. The flow through was collected, concentrated to 500 µl, and subsequently applied to a Superdex 200 increase 10/300 GL (Cytiva) gel filtration column. Fractions containing AF488-labeled C8 were collected and had a AF488 dye to protein label (DOL) of 2.5:1.

### Convertase labeling of bacteria in ΔC5 serum and MAC deposition

Bacteria were grown to mid-log phase as described above, washed and resuspended in RPMI/HSA. Bacteria with an OD600nm ∼0.1 were incubated with 10% C5-depleted serum for 30 minutes, shaking at 37°C. After washing, convertase-labeled bacteria (OD600nm ∼0.05) were incubated with 2.5% serum equivalent C5, C6, C7, C8(-AF488) and C9-AF647 together with Sytox Blue (5 µM) for indicated amounts of time, shaking at 37 °C. At each timepoint, samples were immediately washed and resuspended in RPMI/HSA to OD600nm ∼0.5 for microscopy. For flow cytometry, samples were diluted 10-fold in RPMI/HSA with Sytox Blue (5 µM).

### Antibody-mediated MAC deposition in a purified classical pathway setup

Bacteria were grown to mid-log phase as described above, washed and resuspended in RPMI/HSA. Bacteria with an OD600nm ∼0.1 were incubated with 1 µg/ml anti-GlcNAc IgM and 2.5% serum equivalent C1 for 15 minutes, shaking at 4 °C. Next, C2, C3, C4 and C1-inhibitor were added to a final concentration of 2.5% serum equivalent. After 15 minutes, C5, C6, C7, C8, C9-AF647 were added to a final concentration of 2.5% serum equivalent together with Sytox Blue (5 µM). Bacteria were then incubated for the indicated amounts of time, shaking at 37°C. At each timepoint, samples were immediately washed and resuspended in RPMI/HSA to OD600nm ∼0.5 for microscopy. For flow cytometry, samples were diluted 10-fold in RPMI/HSA with Sytox Blue (5 µM).

### IgM and C3b deposition

Bacteria were grown to mid-log phase as described above, washed and resuspended in RPMI/HSA. To measure anti-GlcNAc deposition, bacteria with an OD600nm ∼0.1 were incubated with 1 µg/ml anti-GlcNAc IgM for 30 minutes, shaking at 4 °C. For IgM detection, bacteria were washed and then incubated with 1.5 µg/ml Goat F(ab’)2 anti-kappa-AF647 (Southern Biotech) for 30 minutes, shaking at 4 °C. To measure C3b deposition, bacteria with an OD600nm ∼0.1 were labeled with convertases by incubation with 10% C5-depleted serum for 30 minutes, shaking at 37°C. Next, bacteria were washed and incubated with 3 µg/ml monoclonal mouse anti-C3b (bH6 (32)) that was randomly labeled with NHS-AF488 (Thermo Fisher Scientific) according to the manufacturer’s protocol. After washing, bacteria were resuspended in RPMI/HSA to OD600nm ∼0.5 for microscopy.

### Widefield microscopy

Agar pads were prepared from 1% low-melting agarose in PBS or RPMI/HSA (for time-lapse imaging). Heated agarose was pipetted on a glass slide and covered with a thick chamber-forming coverslip. If Sytox Blue was measured, Sytox Blue (5 µM) was added to the heated agarose solution before putting it on the glass slide. After solidification of the agarose solution the coverslip was removed. Bacteria at OD600nm ∼0.5 were applied on top of the agar pad and, after drying, covered with a coverslip. Widefield microscopy was performed on a Leica SP5 microscope equipped with a HC PL APO100x/1.40 OIL PH3 objective. AF647, AF488 and Sytox Blue signal were detected using a Y5, GFP/YFP and CFP filter cube, respectively. For time-lapse microscopy, the microscope environment was maintained at 37°C and images were captured every 5 minutes for a period of 2.5 hours.

### Bacterial viability

Convertase-labeled bacteria were incubated with purified C5-C9 and Sytox Blue for different amounts of time as described above. At each timepoint, samples were immediately washed and resuspended in RPMI/HSA. After the assay, samples were serially diluted in PBS and 20 µl droplets were plated in duplicate on LB agar plates. Colonies were counted after overnight incubation at 37 °C to calculate the number of colony forming units (CFUs) per ml in the original sample. Percentage survival was determined by comparison to the sample that had not been incubated with C5-C9 (0-minute timepoint).

### Image analysis

Phase-contrast and fluorescence microscopy images of bacteria were analyzed using a custom Python (version: 3.11.9) script. The script was optimized for high-throughput analysis and supports multiple experimental conditions. It integrates deep learning models for instance segmentation (3-class U-Net (33)) and object classification of bacterial growth stages (Ultralytics YOLO11 (34)).

In our tests, we found that the instance segmentation integrated into YOLO produces artifacts and poor segmentation results for small objects. To address this, a 3-class U-net was trained separately for instance segmentation. In general, using multiple smaller, specialized models provides better performance than a single large, generalized model (35).

### Deep learning models

Our 3-class U-Net was trained to predict the following classes: background, cell membrane (3px wide), cell body. The architecture is based on the residual convolutional neuronal network (adapted from pytorch-3dunet (36)) combined with Spatial and Channel Attention (SCA, (37)) layers with 2,460,562 trainable parameters. We chose for our U-Net 5 layers with 16, 32, 64, 128, 256 channels respectively. Each layer consists of SCA2D, GroupNorm, Convolution2D, ReLu activation with 2x2 max pooling for the down path and transposed convolution upscaling for the up path. A total of 80 images (73 for training and 7 for validation) were manually annotated using Napari (38). Furthermore, full-size grayscale phase-contrast image (1,392 × 1,040 pixels, 0.102 μm pixel size) and the annotated target images were cut into smaller 256 × 256 sub-images (n = 120) with 50% overlap. The model was trained with 27,594 training- and 2,646 validation-image-sets (image augmentation to increase number of image-sets), Adam optimizer (weight decay = 1E-5, learning rate = 5E-4), Tversky and Focal loss using PyTorch (39) and converged in 42 epochs.

Our YOLO model was trained to predict the following classes: dividing, rod shaped, intermediate and defocused bacteria as well as microcolonies. We chose the yolo11m detection model from the Ultralytics YOLO11 implementation with 35,729,296 trainable parameters. A total of 99 images (76 for training and 23 for validation) with 3,358 bounding boxes were manually annotated using a COCO annotator (40) server instance (provided by the University of Applied Sciences Hagenberg). The model was trained with default parameters (batch = 8, image size = 1,024) for 400 epochs.

### Image processing pipeline and fluorescence intensity profile calculation

Each frame of a multidimensional image stack containing a phase-contrast and multiple fluorescence channels (e.g. C9-AF647, …) was successively analyzed. The phase-contrast channel for each frame (timepoint or spatial region) in an experiment was predicted by both deep learning models.

For instance segmentation using our 3-class U-Net model, first the full-size grayscale phase contrast image (1,392 × 1,040 pixels, 0.102 μm pixel size) was cut into smaller 256 × 256 sub-images (N = 120) with 50% overlap. The cut sub-images were then batched (typically batch size of 64 for 6GB of VRAM) and the 3-classes are predicted by the deep learning model. Thereby each pixel was categorized into a class depending on if the pixel contains a cell border, cell body or background. A full-size image (1,392 × 1,040 pixels) was then reconstructed by smooth blending overlapping predicted images (120 sub-images with 256 × 256 pixels). We used a second-order spline window function for blending with 50% overlap (41). Next, cell contours were generated based on the cell body class using OpenCV’s connected component function. Each contour was then rendered to an image again and dilated by exactly 3 pixels. The dilatation was necessary to compensate for the cell border class (which was always annotated to be 3 pixels wide). The reconstructed image (1,392 × 1,040 pixels) represents the instance segmentation mask for the currently processed single cell.

The same phase-contrast image as for the U-Net was then used for object classification using our trained YOLO model. The model predicts multiple objects with a bounding box, classification (dividing, rod shaped, intermediate, defocused bacteria, and microcolonies) and a classification confidence score.

Our algorithm iterates over each cell mask from the segmentation step. Each instance segmented mask of a single cell was then correlated to a YOLO object by highest intersection over union (IoU) value and classification confidence score. Undesired bacterial growth stages (e.g. intermediate, defocused bacteria, and microcolonies) and cell masks intersecting with the image borders were skipped and filtered out. The image was then cropped to the cell mask’s bounding box with an additional small border (typically 20 pixels) around it. Based on this cell mask the long axis of the cell (longest central line or median axis) was then calculated. We offer several algorithms to find the long axis of a cell including skeletonization (42), principal component analysis (43), image moments (44) and elliptical fits. In our experience the image moments gave the most consistent and reliable results of finding the long axis. The long axis was determined using the image moments algorithm by calculating the angle 𝜃 of the long axis from the central moments and cell center (*c_x_, c_y_*) using the spatial moments:

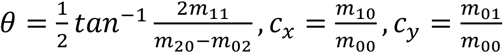

Next, a line with the angle 𝜃 was projected out from the cell center until the border of the cell mask was reached. This long axis line was then fitted using a B-spline of degree 1 (linear fit) for further processing. *N* (typically 21) points were then equidistantly distributed along the long axis line from 5% to 95% of the total length (i.e. line length divided by 100, p_1_ = 5/100, p_2_ = 9.5/100, …, p_20_ = 90.5/100, p_21_ = 95/100). For each point a perpendicular line intersecting with the cell mask was calculated and intensities values for all available fluorescence channels are sampled (typically using sub-pixel values and bilinear interpolation). A fluorescence intensity profile was then calculated by using the maximum pixel values along the sampled perpendicular line. Alternatively, the mean of the pixel values can be used. A comparison of maximum and mean fluorescence intensity profiles using simulated datasets can be found in figure EV3. These fluorescence intensity profiles containing *N* values as well as frame number, class, index, confidence, contour, length and area were saved in a Panda’s data frame for each instance of a segmented cell.

Based on the extracted values, the cells can be further processed to remove outliers or exclude cells with certain conditions (e.g. Sytox-negative). A length threshold can be calculated to filter out misclassified rod-shaped and dividing bacteria. We found that the Otsu threshold algorithm (45) allows reliable separation of the two distributions based on the cell length histogram.

Image thresholding was used to identify cells exhibiting certain conditions based on fluorescence intensities. A reliable threshold was determined by first applying a Gaussian blur (σ = 5 pixels) to the fluorescence channel image, followed by thresholding using the triangular algorithm (46). Cells whose cell mask overlapped with the thresholded image were regarded as positive. Cells lacking fluorescence signals above the calculated threshold within their masks were excluded from further analysis.

A final box plot containing all the fluorescence intensity profiles by class (desired bacterial growth state i.e. Rods, Dividing) was generated after all cells and frames were processed and filtered. For a direct comparison of different fluorescence channels the profiles were normalized to 1.

### Average shapes

The average shape of classified (bacterial growth state: rod shaped and dividing) and filtered bacteria was calculated based on the determined cell mask contours (findContours from OpenCV, which returns a list of vertex positions along the contour) from the image processing pipeline. The contours were then normalized between minus and plus one and rotated so that the long axis of the cell is parallel to the image x-axis. Normalization was facilitated by performing an elliptical fitting the contour positions (fitEllipse from OpenCV, which returns a rotated bounding box). In addition, the number of vertices along the contour of each cell was equalized by interpolation and/or simplification. Next, an average shape is optimized using PyTorch’s ADAM optimizer, where no neuronal network was trained, but the average distance between all vertex positions was minimized. The contour with the lowest Hausdorff distance (47) value was selected for an initial estimate of the average shape and iteratively refined. The mean Hausdorff distance was used as a loss function with a maximum of 1000 iterations, and the optimization was stopped early if the difference between current and previous loss was less than 10^-8^.

### Computation system architecture

Computations for bacteria analysis and development of algorithms were performed on a Dell Precition 3581 notebook (CPU: Intel Core i5-13600H with 12 cores at 2.8GHz, RAM: 32 GB with a speed of 5200 MHz, GPU: NVIDIA RTX A1000 with 6GB VRAM, OS: Window 10 Education operating system 64 bit) and for deep learning on a Dell XPS 8950 workstation (CPU: Intel Core i7-12700k with 12 cores at 3.6GHz, RAM: 64GB DDR5 4600MT/s, GPU: NVidia RTX 3060 with 12GB VRAM, OS: Windows 11 Home operating system 64 bit).

### Flow cytometry analysis

Flow cytometry was performed on a MACSQuant VYB flow cytometer (Miltenyi Biotec). Flow cytometry data were analyzed in FlowJo (v.10). Bacteria were gated based on forward and side scatter. To plot the GeoMFI of C9-AF647 and Sytox Blue in the same graph, log_10_(GeoMFI) values were normalized by setting the minimum and maximum value at 0 and 100, respectively.

### Statistical analysis and graphs

Statistical analysis was performed in GraphPad Prism (v.9) as specified in the figure legends. Graphs were made using GraphPad Prism (v.9) and edited for appearance using Adobe Illustrator.

### Ethics statement

Human blood was isolated after informed consent was obtained from the subject in accordance with the Declaration of Helsinki. Approval was obtained from the medical ethics committee of the UMC Utrecht, the Netherlands.

---

**Figure EV1:**
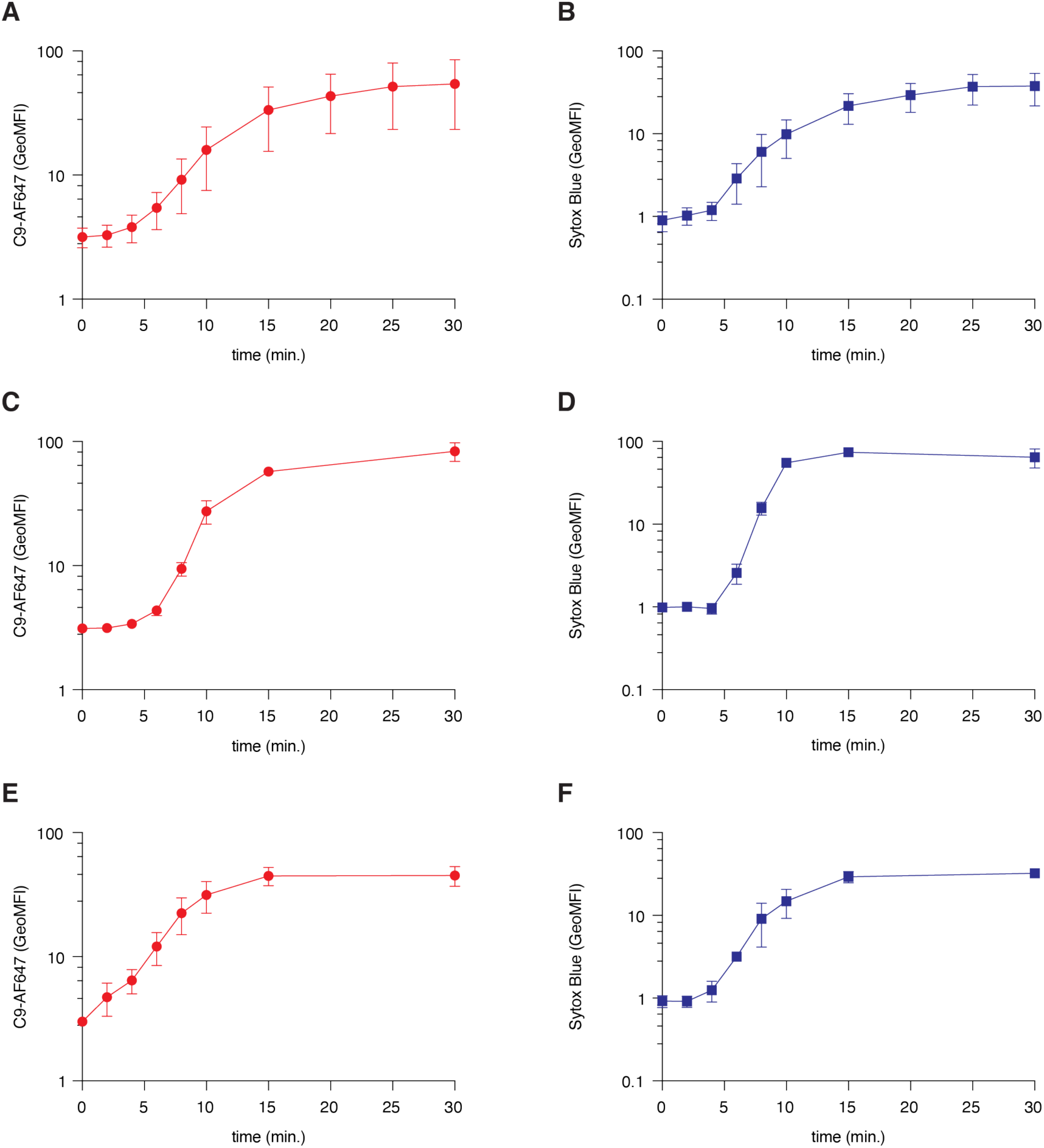
C9-AF647 and Sytox Blue GeoMFI values. (**A, B**) E. coli MG1655 was pre-incubated with C5-depleted serum, washed, and then incubated with purified MAC components for different amounts of time (Fig. 1A). C9-AF647 deposition (A) and inner membrane damage (Sytox Blue) (D) were measured by flow cytometry. Data represent mean ± SD of four independent experiments. (**C, D**) E. coli EC10 was pre-incubated with C5-depleted serum, washed, and then incubated with purified MAC components for different amounts of time (Fig. EV4). C9-AF647 deposition (C) and inner membrane damage (Sytox Blue) (D) were measured by flow cytometry. Data represent mean ± SD of three independent experiments. (**E, F**) MAC deposition was induced on E. coli MG1655 in a purified setup for classical pathway complement activation (Fig. 2A). C9-AF647 deposition (E) and inner membrane damage (Sytox Blue) (F) were measured by flow cytometry. Data represent mean ± SD of three independent experiments.

**Figure EV2:**
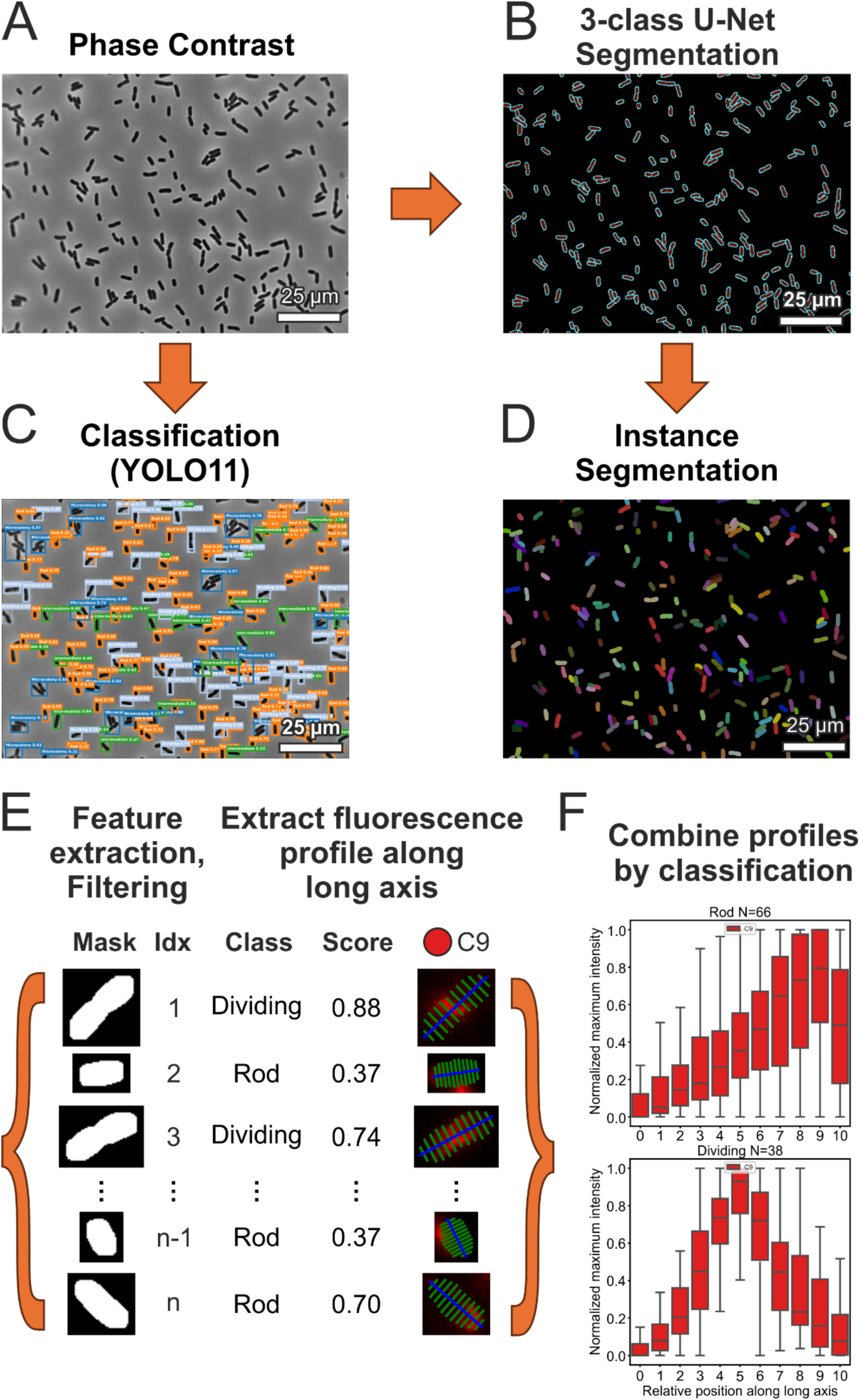
Overview image analysis pipeline of bacteria. (**A**) shows an exemplary phase-contrast microscopy image of Gram-negative bacteria. (**B**) shows semantic segmentation results from a trained 3-class U-Net which predicted bacterial masks for the background, cell membrane and cell body from (A). In (**C**) the object classification results predicted from another custom trained deep learning model (YOLO11) are visualized. Thereby bounding boxes, object classification into bacterial growth stage (microcolonies, dividing, rod shaped, intermediate and defocused bacteria) and classification scores were predicted from the same phase-contrast image as in (A). (**D**) shows false color instance segmentation masks of individual cells based on the 3-class U-Net predictions from (B). Each color represents a single cell with a unique number and mask associated. (**E**) shows the pipeline for processing each instance segmented cell. In this step, cells and their associated mask (cell 1 … n) were further refined. Each cell was then correlated with a matching YOLO result by highest intersection over union value and classification confidence score. Outliers were automatically filtered (i.e. undesired growth stages). Next the geometry of cells is further processed and based on the cell mask the long axis (blue line) of the bacteria is determined by the image momentum algorithm (see Material & Methods chapter “Image processing pipeline and fluorescence intensity profile calculation” for details). The long axis (blue line) was then divided equidistantly into N = 11 points. A fluorescence intensity profile along the long axis was calculated by selecting the maximum pixel value sampled (bilinear interpolation) along the perpendicular lines (green lines). This was repeated for each fluorescent channel and cell. (**F**) visualizes all calculated fluorescence signal profiles along the long axis of individual bacteria split by their growth stage (rod and dividing) in two box plots.

**Figure EV3:**
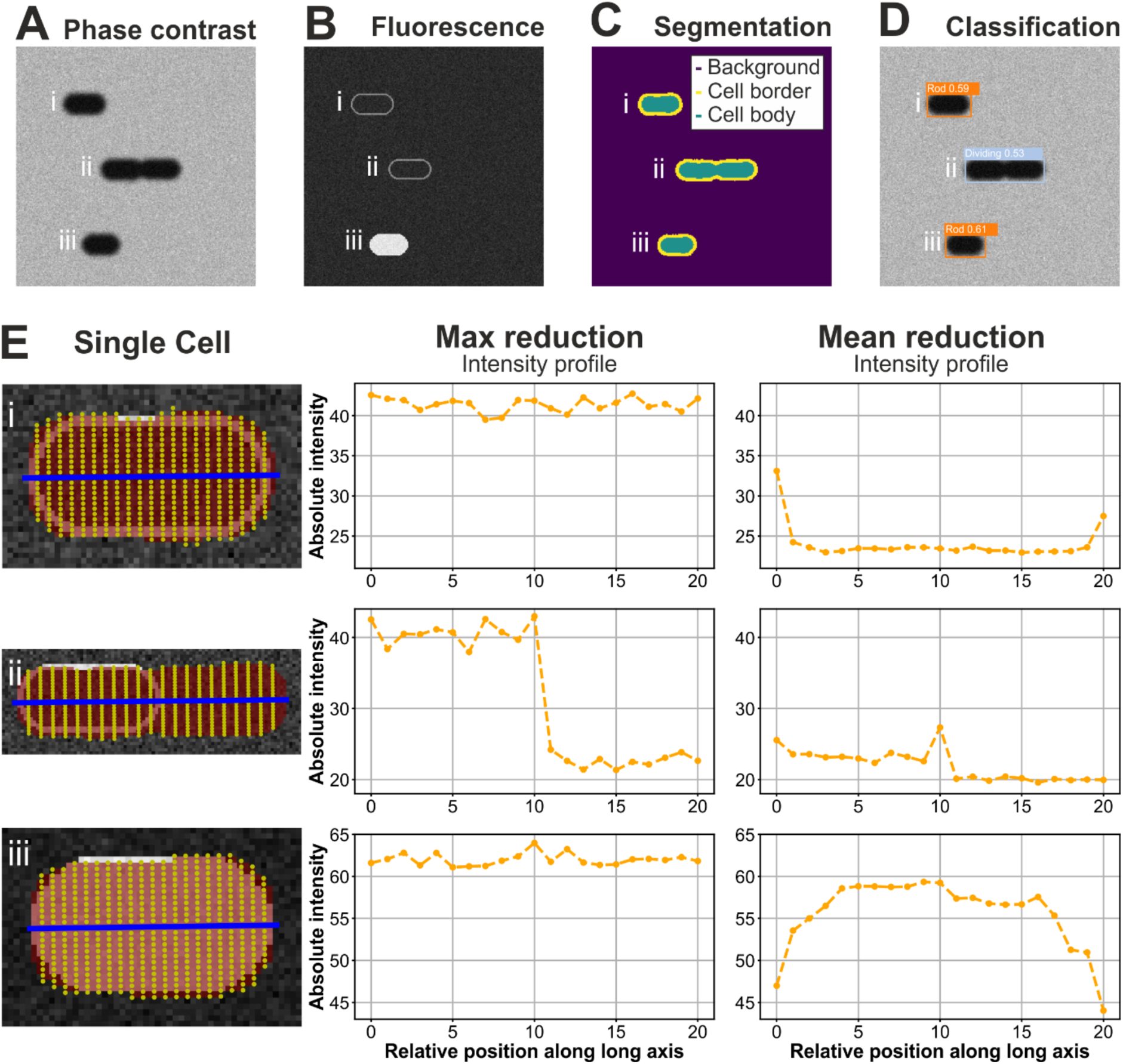
Simulations for the bacteria image analysis pipeline. (**A**) Simulation of phase-contrast microscopy of three bacteria cells at two different growth stages (i+iii: rod shaped, ii: dividing). Noise values (Poisson and Gaussian) were chosen to match the intensity levels of the experiments used in this study (unsigned 8-bit grayscale image with pixels values from 40 to 200). (**B**) Simulation of fluorescent signals of the same three bacteria as in (A) resembling i: signals only in the membrane of a rod-shaped bacterium, ii: signals only in the membrane of a single sub-cell during division, and iii: signals in the whole cell with the same intensity. The pixel intensity values were again chosen to match experimental conditions (unsigned 8-bit grayscale image with pixels values from 10 to 66). (**C**) Segmentation of the simulated phase contrast image from (A) using our trained deep neuronal network U-Net with 3 classes. (**D**) YOLO object detection and classification of the simulated phase contrast image from (A) using our trained YOLO11 deep neuronal network. (**E**) Application of our image analysis pipeline to all three simulated bacteria cells (one cell in each row). The first column shows the simulated fluorescence signal from (B) in the background of a single cells, in red the determined instance segmentation mask from (C), in blue the calculated long axis using the image momentum algorithm, and each yellow dot represents sampled pixels (bilinear interpolation) along a perpendicular line (to the long axis line) at equidistant points along the long axis line (N = 21). The second and third column show the calculated fluorescence intensity profile, where for each point along the long axis line (blue), the intensity values along the perpendicular line (yellow dots) were combined (reduced) by either the maximum value (second column) or the mean value (third column).

**Figure EV4:**
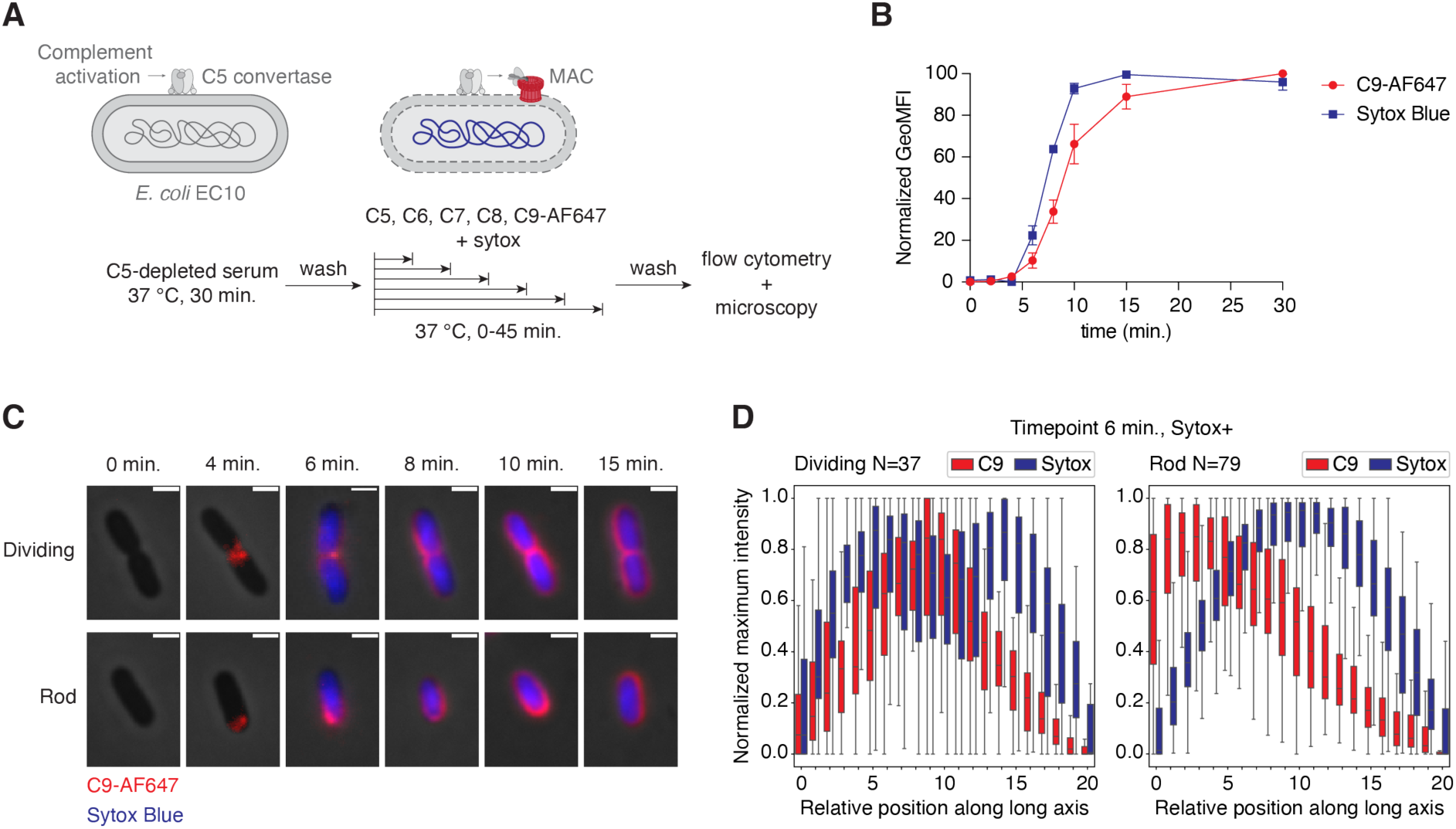
MAC localization on a clinical E. coli isolate. (**A**) Schematic overview of the experimental setup used to study MAC deposition and inner membrane in time. E. coli EC10 was pre-incubated with C5-depleted serum, washed, and then incubated with purified MAC components for different amounts of time. (**B**) C9-AF647 deposition and inner membrane damage (Sytox Blue) in time, measured by flow cytometry. GeoMFI values were normalized by calculating the percentage of the maximum value after log-transformation. Data represent mean ± SD of three independent experiments. (**C**) Phase-contrast and overlayed fluorescence images of selected bacteria at different timepoints. C9-AF647 is shown in red and Sytox in blue. Individual bacteria have been chosen to reflect an average bacterium in the sample and are representative for three independent experiments. Scale bars, 2 µm. (**D**) Normalized average C9 and Sytox distribution along the long axis of dividing and rod-shaped bacteria after 6 minutes incubation with C5-C9. Data analysis was performed on 14 images of one independent experiment.

**Figure EV5:**
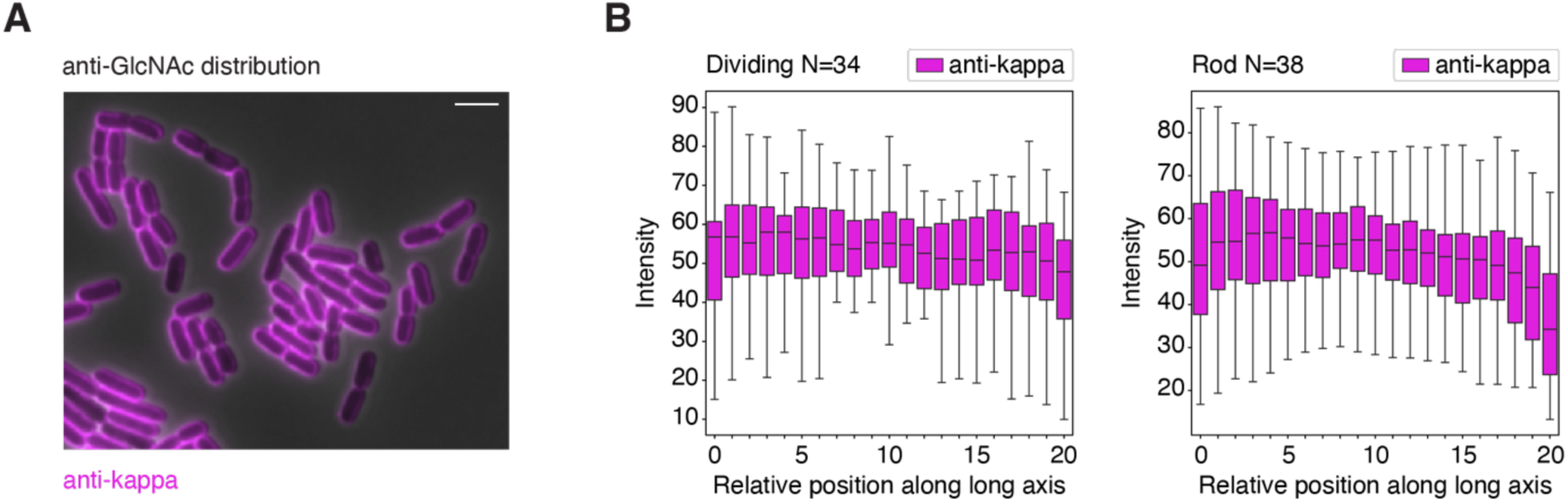
anti-GlcNAc IgM distribution on E. coli. (**A**) Bacteria were incubated with 1 µg/ml anti-GlcNAc IgM followed by staining with a fluorescently labeled anti-kappa F(ab’)2 fragment. Scale bar, 5 µm. (**C**) Average anti-GlcNAc IgM distribution along the long axis of dividing and rod-shaped bacteria. Data analysis was performed on ten images of one independent experiment.

**Figure EV6:**
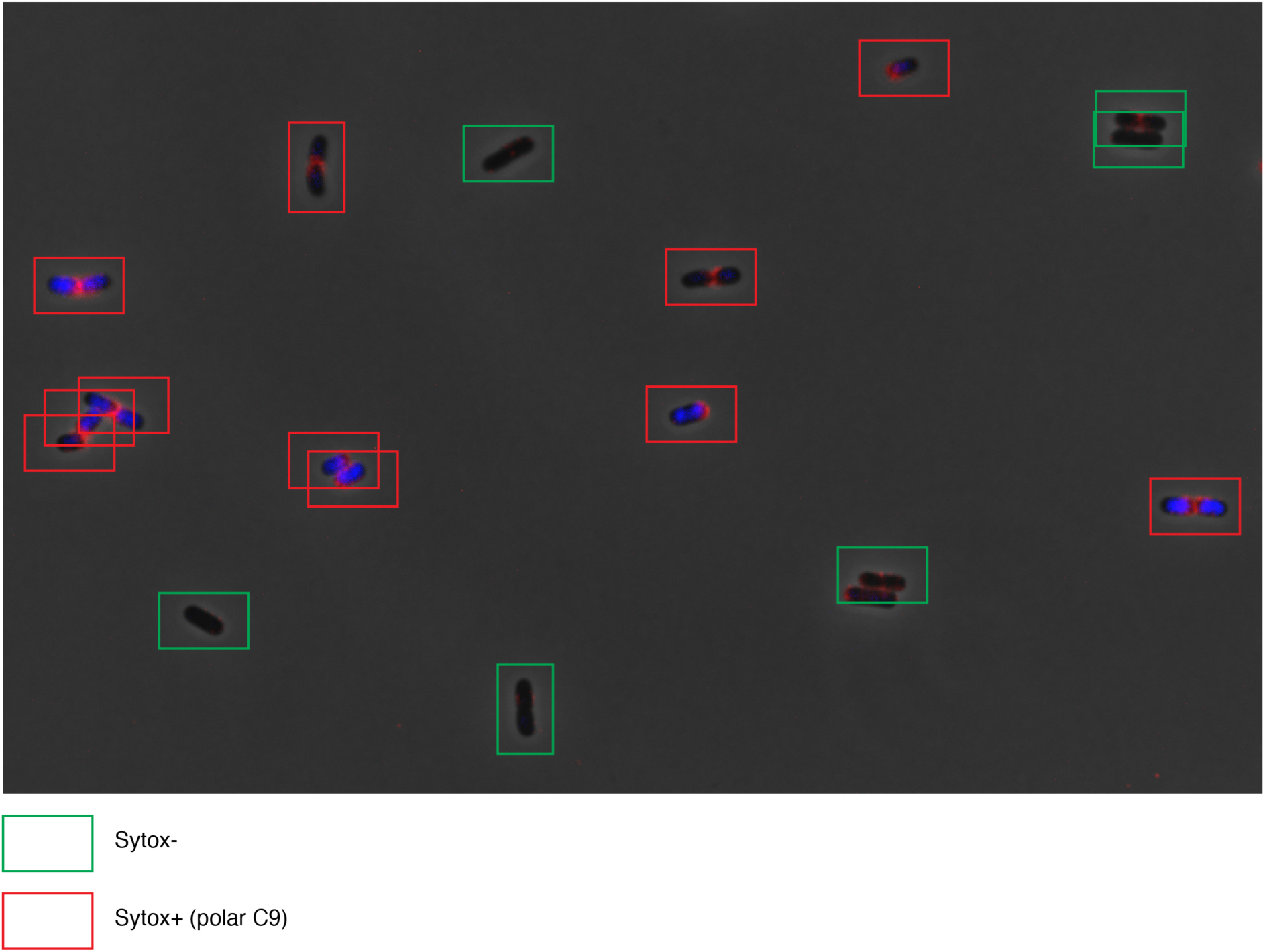
Classification of bacteria at the start of time-lapse imaging. In this example, E. coli MG1655 had been pre-incubated with C5-depleted serum, washed, and then incubated with purified MAC components for 6 minutes (Fig. 4). Bacteria were manually classified based on Sytox-positivity and MAC localization. Sytox-negative bacteria are annotated with green boxes and Sytox-positive bacteria with polar MAC localization with red boxes.

